# HIV remodels germinal centers through T follicular helper cell loss

**DOI:** 10.64898/2026.10.08.757815

**Authors:** Derek R. Jacobs, Faiaz S. Abdool, Lily F. Pohlenz, Danielle P. Curtis, Raman K. Gill, John Peyton Bush, Michael S. Martin, Julia Toman, Warren Kuhn, Sarena Maistry, Ashley Levy, Ronald S. Veazey, Meagan Watkins, Jay Rappaport, J. Zachary Porterfield

## Abstract

T follicular helper cells (Tfh) within germinal centers are among the most permissive cells for HIV infection and serve as a critical part of the viral reservoir. Given the key role of Tfh in affinity maturation and the production of plasma cells and memory B cells, an understanding of this critical viral reservoir is fundamental not only for cure strategies but also for understanding immunodeficiency in people living with HIV (PLWH). Using rare clinical specimens from PLWH, we demonstrate that HIV causes both a change in the cellular composition of germinal centers and changes in their overall architecture in ways that are consistent with the immunodeficiency seen in PLWH, independent of their CD4 count. We note a significant decrease in the number of GC Tfh cells and an increase in the number of follicular CD8 T cells in lymphoid tissues obtained from PLWH. Our data further show that GC size is decreased both by HIV and with increased age. Single-cell RNA sequencing data demonstrate that HIV acutely decreases the expression of Tfh B-cell support pathways and disrupts Tfh expression of cell trafficking receptors. Additionally, in a lymphoid organoid model of acute infection that forms GC-like structures, these findings are recapitulated, suggesting that New Approach Methodologies could be germane to understanding the impact of HIV on the reservoir and immunodeficiency. This study describes previously undocumented remodeling of the germinal center niche in PLWH, providing insight into the immune microenvironment of a key HIV reservoir that may help inform future cure strategies.

**Importance:** Human immunodeficiency virus (HIV) is a chronic viral infection that affects more than 40 million people worldwide. Even with antiretroviral therapy, people living with HIV remain at increased risk for a variety of comorbidities and exhibit persistent immune dysfunction that may worsen with age. Understanding how HIV impacts key immune structures such as germinal centers is important for preventing or reversing these effects. This study reveals changes in germinal center composition and cellular function, including decreased T follicular helper cells and germinal center size, that may contribute to persistent immune dysfunction in people living with HIV. Our findings provide new insight into how HIV alters the lymphoid tissue microenvironment despite antiretroviral therapy and establish a lymphoid organoid model for investigating these changes, which may help identify new strategies to reduce HIV-associated morbidity.

## Introduction

HIV remains a serious infectious disease, causing significant morbidity and mortality, with an estimated 40.8 million people living with HIV (PLWH) globally (1). Although antiretroviral treatments (ART) are highly effective at achieving an undetectable viral load in the blood, HIV remains latent during ART and can re-emerge if treatment is interrupted (2, 3). This latent viral reservoir prevents patients from achieving sustained ART-free remission (2). Within secondary lymphoid tissues, the B-cell follicles and the germinal centers (GCs) within them constitute one of the most significant HIV reservoirs, especially in patients on suppressive ART (4). GCs are primarily composed of B cells, T follicular helper cells (Tfh), and follicular dendritic cells (fDCs)(5). It is well described that the GC niche is a highly orchestrated and tightly regulated system with a defined morphology and organization that regulates B cell responses (6–8).

Tfh cells are known to be a predominant site of HIV replication and persistence both on and off ART (4, 9–12). Tfh cells orchestrate immune function by supporting GC development and by mediating the selection and survival of antigen-specific B cells during GC reactions (5, 13, 14). These functions are essential for the development of memory B cells and plasma cells, which generate highly specific antibodies and are critical to adaptive immunity (15). In HIV, dysfunction in memory B cells is associated with poor immune recovery after starting ART (16). Prolonged Tfh activation and high functionality during HIV infection are also associated with improved development of neutralizing antibodies, further highlighting their importance in anti-HIV immune responses (15). It is well characterized that during HIV infection, Tfh-B-cell signaling is disrupted (17, 18). Although some studies performed using flow cytometry techniques on cells isolated from lymphoid tissue have noted an expansion in Tfh cell number, a systematic analysis of Tfh cells in the GC microenvironment of human tissue in the setting of HIV has not been performed (9, 19). Furthermore, some studies have identified changes in Tfh phenotype during HIV infection, but a deeper analysis of how this phenotype is related to the reservoir remains unexplored (19, 20). Of note, in a non-human primate (NHP) macaque model of untreated SIV infection, it has been demonstrated that splenic Tfh cells decrease following 180 days of infection with SIVmac251 (21).

It is well documented that PLWH are at a much higher risk of developing a variety of comorbidities, including cardiovascular disease, cancer, and diabetes mellitus (22). More broadly, Tfh are implicated in a variety of disease states, including infection, autoimmunity, and systemic pathologies such as atherosclerosis and cancer (14). Whether the impacts of HIV on Tfh and GCs can be linked more broadly to the development of these comorbidities is an open question. Overall, the GC reservoir is widely acknowledged as a critical barrier to establishing a durable HIV cure, and yet the response of this unique immune microenvironment to HIV infection has not been fully described in human tissues. Studying these changes necessitates access to difficult-to-acquire tissue samples, which requires a robust collaboration between communities, clinicians, and research teams. Our research team and collaborators have established multiple such cohorts, which we leverage here to describe an in-depth analysis of alterations in GC cell demographics, shape, size, cellular content, and signaling mechanisms that explores HIV’s impacts on these questions.

## Results

### Cohorts Developed for These Analyses

Critical to the assessment of changes in germinal center structure and function, is the acquisition of lymphoid tissue from PLWH and people living without HIV (PLWOH). For these analyses, we have developed or contributed significantly to multiple tissue biobanks. These analyses are derived from samples from 21 PLWH, 6 PLWOH at the African Health Research Institute (AHRI) in KwaZulu-Natal, South Africa, and 34 PLWOH from the University of South Florida in Tampa, Florida. Notably, we selected biobanked samples from a relatively broad range of ages and HIV statuses to support generalizability, acknowledging the need for further subgroup analysis.

Overall, the average age of study participants was 20.4 years and 71% were female. Among the PLWH, the average CD4 count was 518 cells/mm^3^ and 60% of PLWH had a viral load below the limit of detection at the time of sample acquisition, with a median viral load of 42,068 for those with detectable virus. Amongst the South African cohort, the average age of PLWH was 28.7 years (standard deviation of 12.0 years), while the average age of PLWOH was 31.8 years (standard deviation of 9.9), with no statistical difference in age between these groups. For samples obtained from the Tampa cohort, all of which were PLWOH, the average age of participants was 14.7 years (standard deviation of 11.5 years). The difference in ages between the Tampa participants and the South African participants, both PLWH and PLWOH, were not statistically significant. Given limited sample availability from the South African cohort, complete age matching between PLWH and PLWOH from the South African cohort alone was not possible. Since no statistical difference in age existed between the Tampa and South African cohorts, we selected the six available samples from the South African PLWOH cohort and combined this cohort with 11 adult participants from the Tampa cohort of PLWOH to allow for complete representative age matching of South African PLWH cohort. Although only two child samples from the South African PLWH cohort were available, to evaluate age related effects in younger patients, 18 child samples from the Tampa PWLOH cohort with an average age of 7.6 years (standard deviation of 3.3 years) were included.

### PLWH have a reduced GC Tfh percentage and a decreased Tfh density

We examined changes in GC Tfh cells related to HIV through IHC staining for PD1 in formalin-fixed, paraffin-embedded (FFPE) samples. While PD1 expression is classically associated with T cell exhaustion, within the context of the GC, it is a canonical Tfh marker given that Tfh express PD1 at high levels and there is a paucity of other T cell subsets within GCs (23). Notably, IHC staining for PD1 demonstrated light zone/dark zone polarization classically associated with Tfh cell distribution across the light zone in GCs (Fig 1A). We observed a significant decrease in GC Tfh cells, with Tfh cells comprising 16.1% of all GC cells in PLWH versus 24.6% in PLWOH (Fig 1B). The percentage of GC Tfh cells did not correlate strongly with patient CD4 count or viral load (Fig 1C, Supplemental Fig 1A). Due to increasing evidence of immune dysfunction with advanced age, which may be exacerbated in PLWH, we sought to evaluate the effects of age on GC Tfh cells (24, 25). A slight trend toward decreased GC Tfh cells with age was seen in PLWOH; however, this was exacerbated in PLWH, although the trend was not statistically significant (p=0.07) (Fig 1D and E). We sought to assess whether the decrease in Tfh number impacted density and distance between Tfh cells. PLWH had a decreased Tfh cell density of 0.031 Tfh/µm^2^ and increased mean distance from Tfh to Tfh of 3.45µm, whereas PLWOH had a Tfh density of 0.045 Tfh/µm^2^ and a Tfh-Tfh distance of 2.72µm on average (Fig 1F and G).

**Fig 1.**
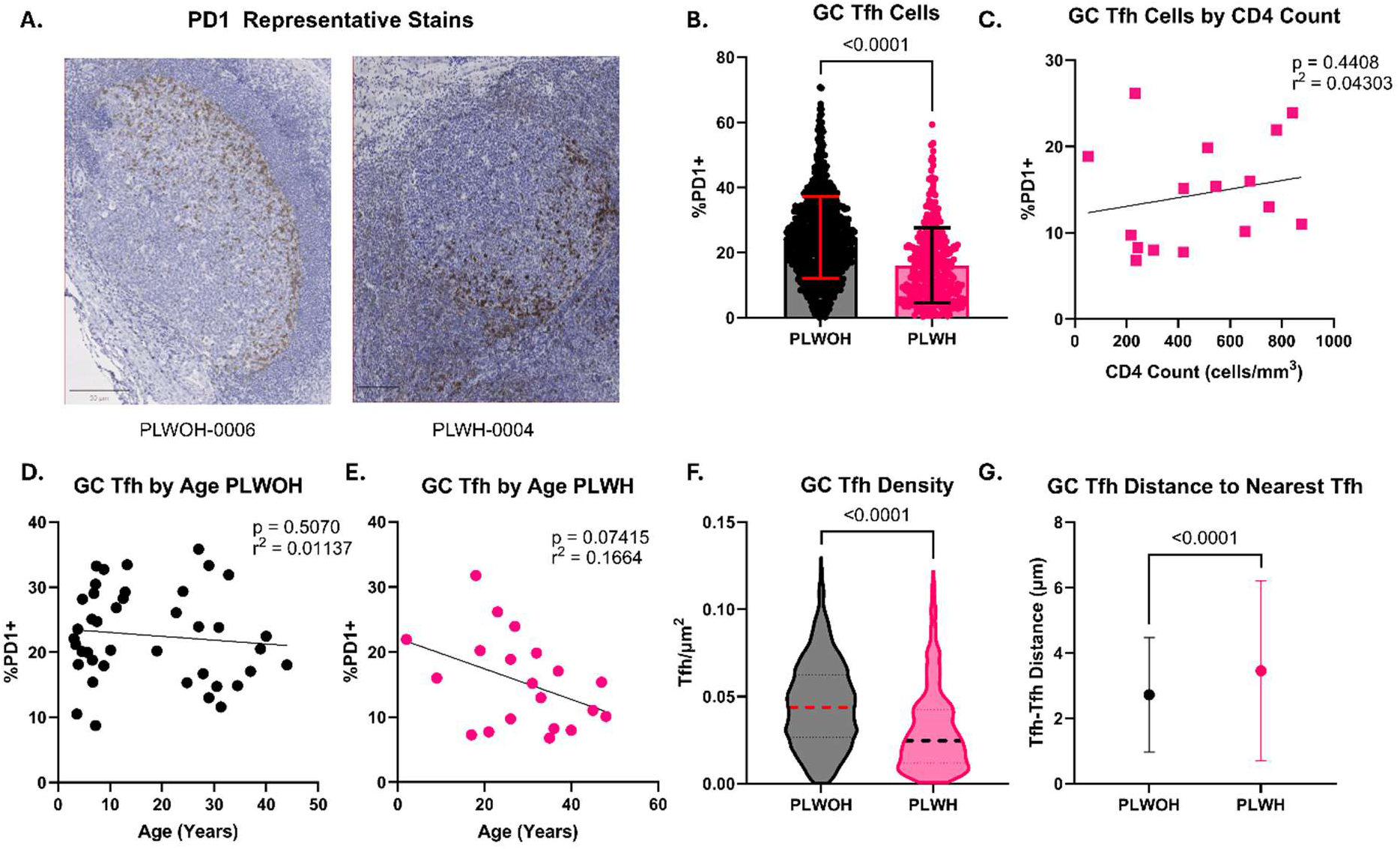
GC Tfh are decreased in number and density in PLWH. **A.** Representative PD1 IHC stains, representing GC Tfh cells. **B.** Comparison of percent GC PD1+ (Tfh) cells in 39 PLWOH (black) vs 20 PLWH (pink). Each dot represents an individual GC. **C.** Linear regression of percentage of Tfh cells and CD4 count for samples from PLWH (n=16 patients), with each data point representing the average percent GC Tfh for that patient. **D.** Linear regression of percentage of Tfh and patient age in PLWOH (n=39 patients). **E.** Linear regression of percentage of Tfh and patient age in PLWH only (n=20 patients). **F.** Comparison of GC Tfh density (Tfh/µm^2^) between PLWOH (n=39) and PLWH (n=20). **G.** Comparison of GC Tfh distance to nearest Tfh (µm) between PLWOH (n=39) and PLWH (n=20).

### GC size decreases with HIV and age

Studies of GC morphology prior to the ART era noted an increase in GC size and shape eccentricity in post-mortem cohorts of PLWH; thus, we sought to examine these features in our sample set. We noted a significant decrease in GC size in PLWH, with an average GC cross-sectional size of 10,966µm^2^, while PLWOH GCs were 18,398µm^2^ on average (Fig 2A).

**Fig 2.**
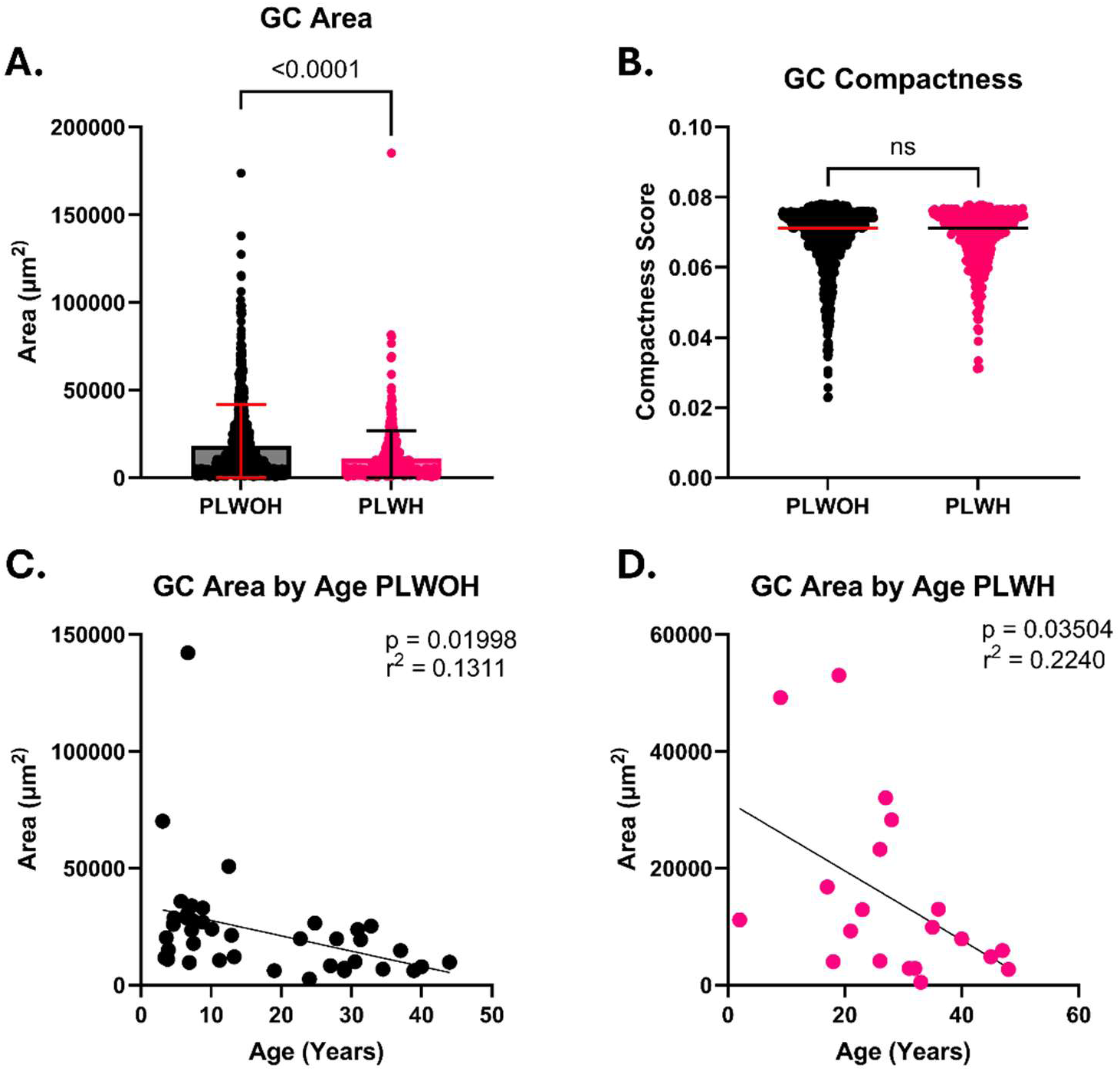
HIV and age cause decreased GC size. **A.** A comparison of GC area (µm^2^) between PLWOH (n=39) and PLWH (n=20). **B.** A comparison of GC compactness (Area/Perimeter^2^) between PLWOH (n=39) and PLWH (n=20). **C.** Linear regression of GC area and patient age in PLWOH only (n=39). Each dot represents the mean GC size for that patient. **D.** Linear regression of GC area and patient age in PLWH only (n=20). Each dot represents the mean GC size for that patient. Note: ns = p>0.05.

Germinal center shape, measured via a compactness score of area/perimeter^2^ (a unitless means of measurement in which perfect circular shapes are maximally compact), was not significantly altered in PLWH (Fig 2B). Given the association of advanced age and immunodeficiency, we sought to determine whether age correlated with GC size. In PLWOH, we observed a modest but significant trend toward decreased GC size with increasing age, r^2^=0.13 (Fig 2C). This trend was recapitulated in PLWH, but notably, the correlation was stronger, r^2^=0.22 (Fig 2D).

### PLWH have increased GC CD8 T cells and decreased GC macrophages

We then sought to determine how other cell types and viral proteins changed in the GC environment in PLWH. Our immunohistochemistry data demonstrate a strong correlation between the percentage of GC cells staining positive for HIV p24 capsid protein and the viral load of the patient, r^2^=0.73 (Fig 3A and B). After p24 staining, we moved on to evaluating changes in other HIV-relevant cell types. While the exact role of macrophages as a functional HIV reservoir remains debated, we evaluated the number of GC macrophages to examine changes in potential reservoir cells. These data show that macrophages are relatively rare in GCs, with a mean of 2.3% of GC cells staining CD68+ in PLWOH. PLWH had a small but statistically significant decrease to a mean of 2.1% (Fig 3C). GC macrophages did not correlate with age, CD4 count, or viral load (Supplemental Fig 1B, C, & D). CD8 T cells, overall rare in GCs, displayed a 6-fold increase in prevalence within GCs, from 1.4% of all GC cells in PLWOH to 9.6% in PLWH (Fig 3D). The percentage of GC CD8 T cells also had a statistically significant correlation with patient viral load, but not patient CD4 count (Fig 3E and F). To investigate whether the increased presence of CD8 T cells in GCs was associated with the observed decrease in GC Tfh cells and macrophages in PLWH, we started by using multicolor immunofluorescence to evaluate the spatial relationship between these cell types (Fig 5A and Supplemental Fig 3). In PLWOH, the mean distance was 78.5µm from a GC Tfh cell (PD1+) to a cell expressing CD8.

**Fig 3.**
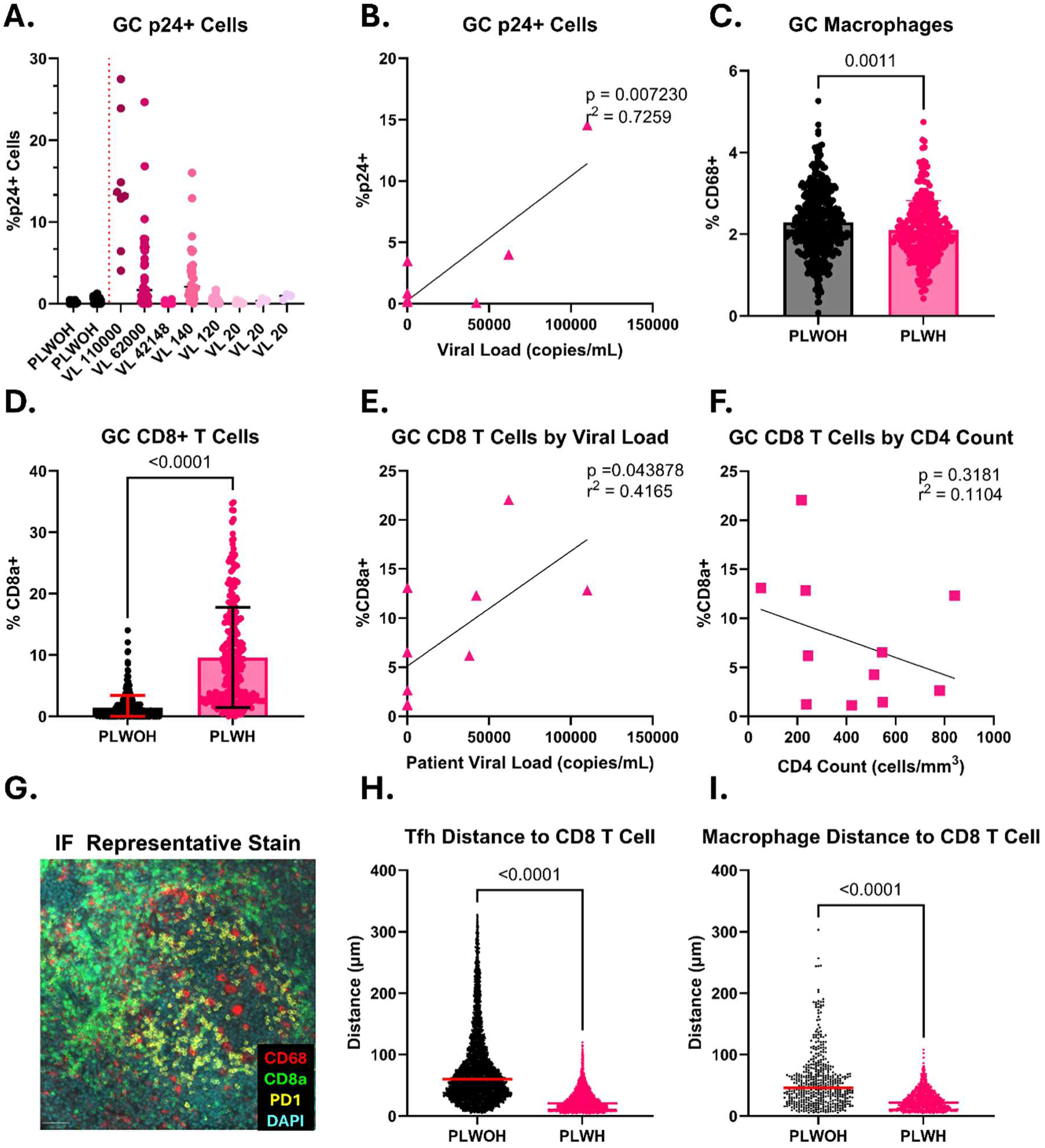
Changes in germinal center HIV protein and CD8 T cells in the setting of HIV. **A.** Patient-by-patient breakdown of GC cells with positive p24 IHC staining for study participants with viremia as well as two PLWOH and three PLWH with suppressed viral load. Each dot represents an individual germinal center. **B.** Linear regression of percent of p24+ GC cells as a function of patient viral load, with each dot representing the mean for that patient. **C.** Comparison of percent GC CD68+ (macrophage) cells in PLWOH (n=13) and PLWH (n=13). Each dot represents an individual GC. **D.** Comparison of percent GC CD8a+ (CD8 T cells) cells in PLWOH (n=16) vs PLWH (n=13). Each dot represents an individual GC. **E.** Linear regression of percent CD8a+ GC cells versus patient viral load. Each dot represents the mean for that patient. **F.** Linear regression of percent CD8a+ GC cells and CD4 count. Each dot represents the mean for that patient. **G.** A representative image of immunofluorescence staining for CD68, CD8a, and PD1 serving as a markers for germinal center macrophages, CD8 T cells, and Tfh cells respectively. **H.** Quantification of distance between Tfh cells (PD1+) and the nearest cell expressing CD8a in PLWOH (n=2) and PLWH (n=2). Each dot represents a single cell-cell distance. **I.** Quantification of distance between macrophage cells (CD68+) and the nearest cell expressing CD8a in PLWOH (n=2) and PLWH (n=2). Each dot represents a single cell-cell distance. Note: VL = viral load.

This distance decreased to 35.9µm in PLWH (Fig 5B). Likewise, the macrophage-CD8 T cell mean cell-cell distance in PLWOH was 57.6µm and decreased to 25.5µm in PLWH (Fig 5C).

### Acute ex-vivo HIV infection reduces Tfh support for B cells and disrupts Tfh chemokine related gene expression

To further explore the impact of HIV infection on Tfh cells within GCs, we analyzed publicly available single-cell sequencing data from ex vivo HIV-infected tonsil tissue (26). The study by Freeman et al. focused on the activation of oxidative phosphorylation in CD4 T cells infected with HIV and used tonsil explants from 7 donors that were exposed to HIV and harvested for single-cell RNA sequencing (scRNAseq) (Supplemental Fig 4A-C). Re-examining this dataset through the lens of changes to viral reservoirs and Tfh expression signatures, we again identified a trend toward decreased Tfh cells and an expansion of CD8 T cells that was consistent with what was observed in the GCs of chronically infected PLWH (Fig 4A-C, Supplemental Fig 4C). Applying a gene ontology (GO) biological process analysis to the differentially expressed genes in the Tfh cell subcluster, we recapitulated many of the key findings that the original authors observed, chiefly substantial metabolic changes in HIV-exposed Tfh cells (26). However, this analysis also revealed stark differences in the enrichment of pathways relating to antiviral responses, B cell support, and lymphocyte trafficking between HIV-exposed and HIV-naïve Tfh cells. HIV-exposed Tfh cells had low enrichment of pathways related to viral defense and antiviral response relative to unexposed controls, a finding supporting HIV’s well-documented ability to block and evade host immune responses (Fig 4D) (27, 28). Lower expression of pathways related to leukocyte activation and decreased expression of critical chemokine receptors such as CXCR4 and CCR7 were observed in HIV-exposed Tfh cells (Fig 4D and E). Pathways and genes related to B cell activation, differentiation, and support were also noted to be lower in HIV-exposed Tfh cells (Fig 4D and E).

**Fig 4.**
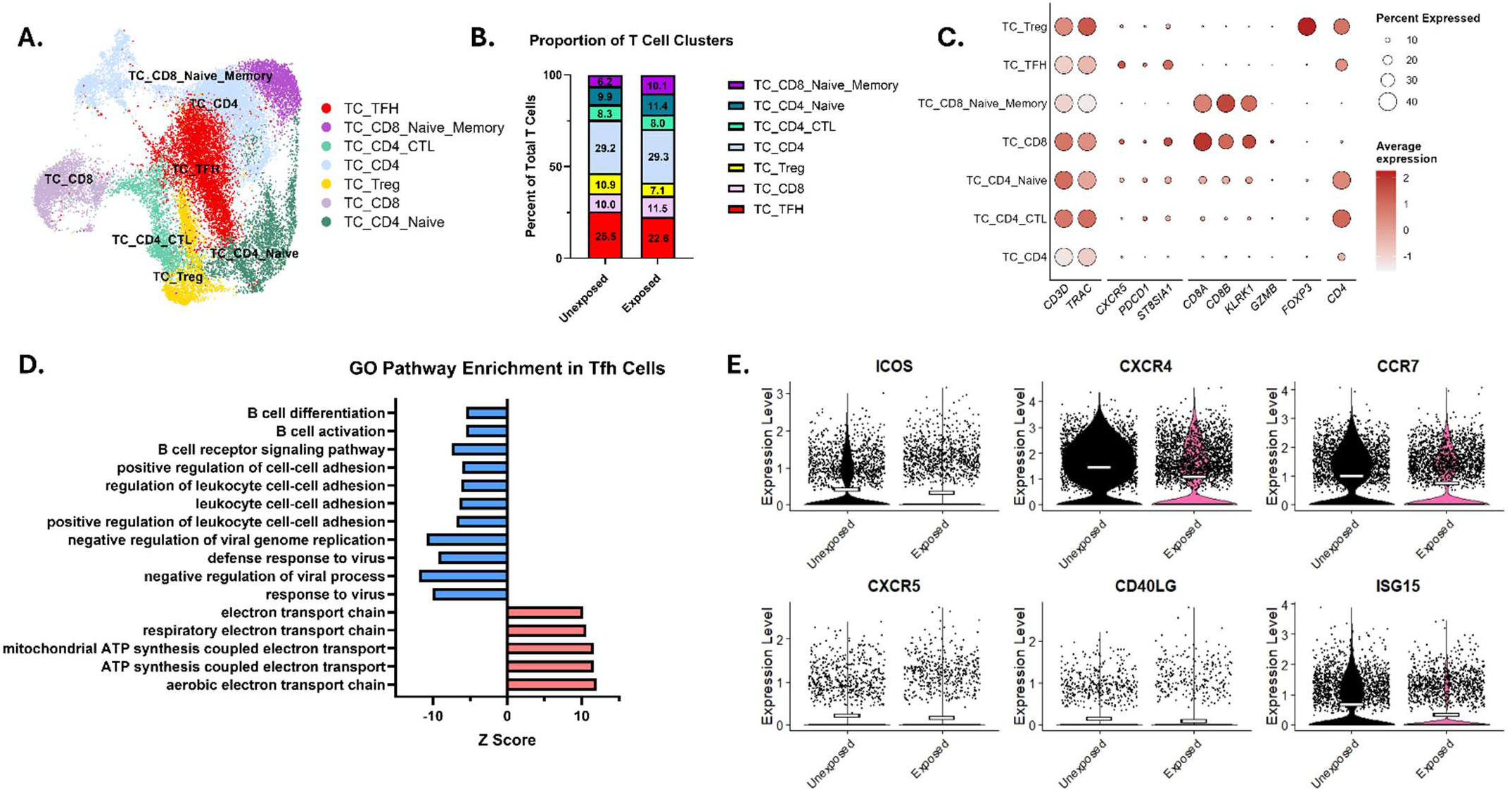
Tfh cells have disrupted B cell support and cell-cell adhesion pathways in HIV exposed tonsil explants. **A.** Annotated UMAP of T cell subset from whole tonsil from publicly available data (k = 26,815 T cells). **B.** Proportion of T cell subtypes in HIV exposed vs unexposed conditions (k = 15,969 HIV exposed cells, k = 10,849 unexposed cells). **C.** Dot plot of T cell markers used to assign cluster identities. **D.** Gene ontology analysis of pathways upregulated (pink) and downregulated (blue) in HIV exposed vs unexposed Tfh cells were performed separately. **D.** Violin plot of differentially expressed genes in exposed vs unexposed Tfh cells, mean expression level marked by the white bar.

### A CXCR4-tropic HIV lineage causes Tfh cell depletion in an organoid model of acute infection

To further explore these findings, we exposed our lymphoid organoid model to multiple virus lineages, including HIV NL4-3 (a lab-adapted, subtype B, X4-tropic variant) and NL4-3 ADA-GFP, in which the envelope has been substituted with envelope from the ADA strain (an R5, macrophage-tropic variant) and a GFP tag has been added, to determine if acute HIV infection in vitro recapitulated the changes in GCs observed in the tissue samples derived from PLWH. In culture, these organoids form GC-like structures that can generate antibody responses that undergo affinity maturation and mount antibody-mediated responses to novel antigens. We initially assessed the total area covered by these GC-like structures as well as their average size.

In our HIV-infected organoids at day 1 and day 4, total GC-like surface area and average GC- like size were not significantly different between conditions (Supplemental Fig 5A and B). At day 7, however, there was a clear trend in both HIV conditions toward reduced total GC-like surface area that did not achieve statistical significance (Fig 5B and C). Mean GC-like size also trended toward being smaller in the NL4-3-infected organoids, but this did not achieve statistical significance (Fig 5D).

**Fig 5.**
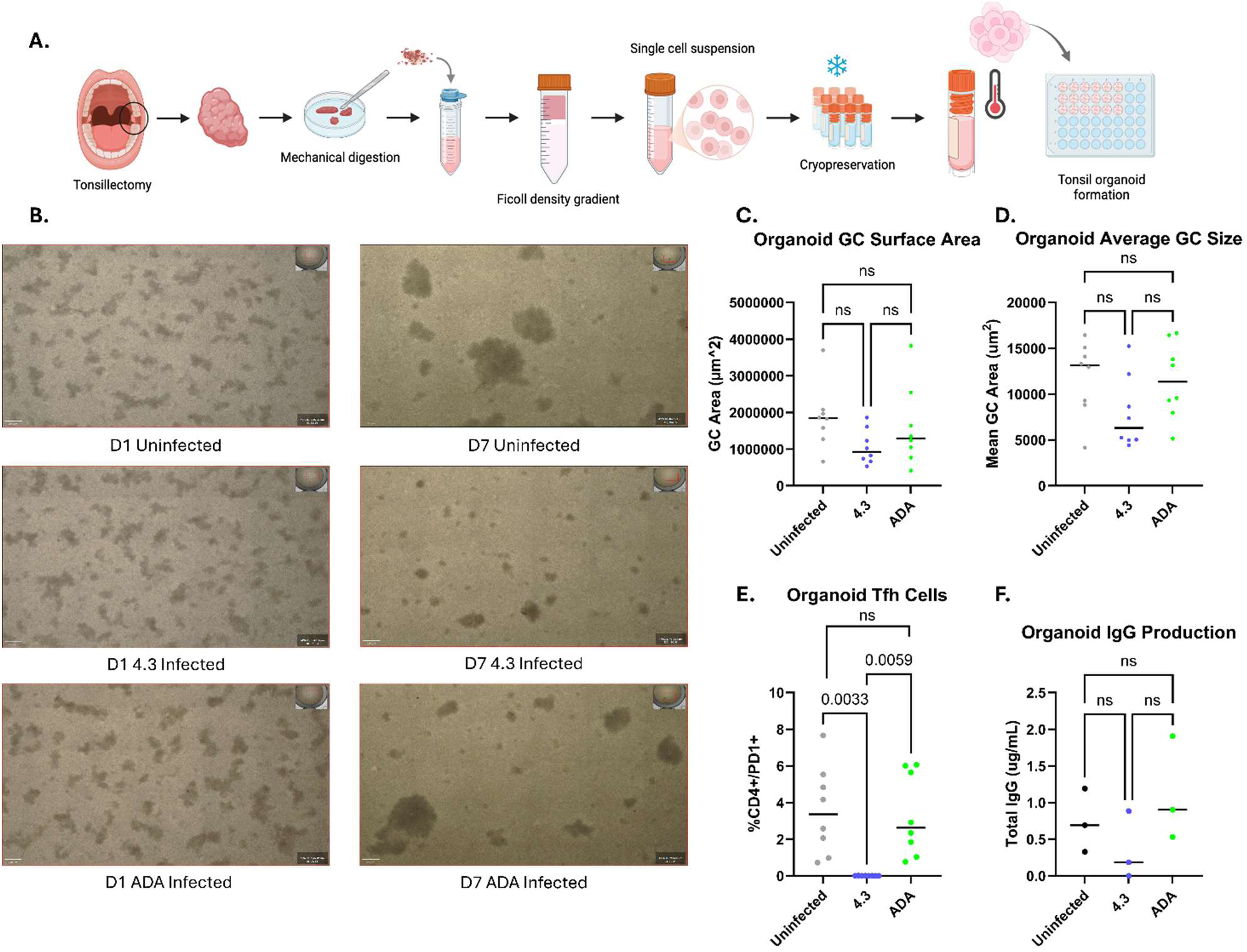
HIV decreases GC-like surface area in tonsil immune organoids. **A.** A schematic overview of the tonsil organoid protocol. **B.** Representative images of organoids in each group taken on day 1 and day 7 showing reorganization into GC-like structures. Magnification level is the same in each image. **C.** A quantification of the day 7 total GC-like organoid surface area in each condition (n = 8 patients). **D.** Quantification of mean GC-like organoid size (n = 8 patients). **E.** A comparison of the Tfh cells in each organoid condition determined by flow cytometry. Values represent the percentage of CD4+/PD1+ cells of all living organoid cells (n = 8 patients). **F.** Total IgG from immune organoid media collected on day 10 quantified by ELISA. n = 3 patients. Note: ns = p>0.05.

Notably, in the NL4-3-infected organoids, there was a significant decrease in Tfh cells, with Tfh cells being almost completely absent, while the uninfected and ADA wells contained roughly 3% Tfh cells (Fig 5E). In order to assess GC function, we measured the total IgG antibody amount in the organoid media on day 7 and day 10. While B cells are responsible for producing IgG, Tfh cells are critical for affinity maturation through their activation and support of B cells. We noted that at day 7, there was no difference in IgG production between conditions. At day 10, there was a trend toward decreased IgG production in the NL4-3-infected organoids that did not achieve statistical significance (Supplemental Fig 5C, Fig 5F).

### Adenoviral vaccine that primes T cell-mediated response is associated with increased GC Tfh in a macaque model of SIV infection

In order to evaluate the in vivo impacts of a novel anti-HIV preventative intervention on the GC microenvironment, we assessed non-human primate tissue samples to determine if anti-SIV vaccination impacts GC Tfh cell proportion during SIV infection. We used tissues from 8 rhesus macaques that were exposed to SIVmac251. Prior to SIV exposure, 6 primates received a T cell-priming adenoviral vaccine, while 2 received an empty vector vaccine. Because NHPs have anatomical differences in palantine tonsils, we performed IHC on other lymphoid tissues, including spleens and mediastinal, axillary, and mesenteric lymph nodes. We determined that the vaccinated NHPs had GCs containing an average of 25.8% Tfh cells, while the sham-vaccinated NHPs had a mean of 22.1% (Fig 6A). GC macrophages were not impacted by the type of vaccine given (Fig 6B).

**Fig 6.**
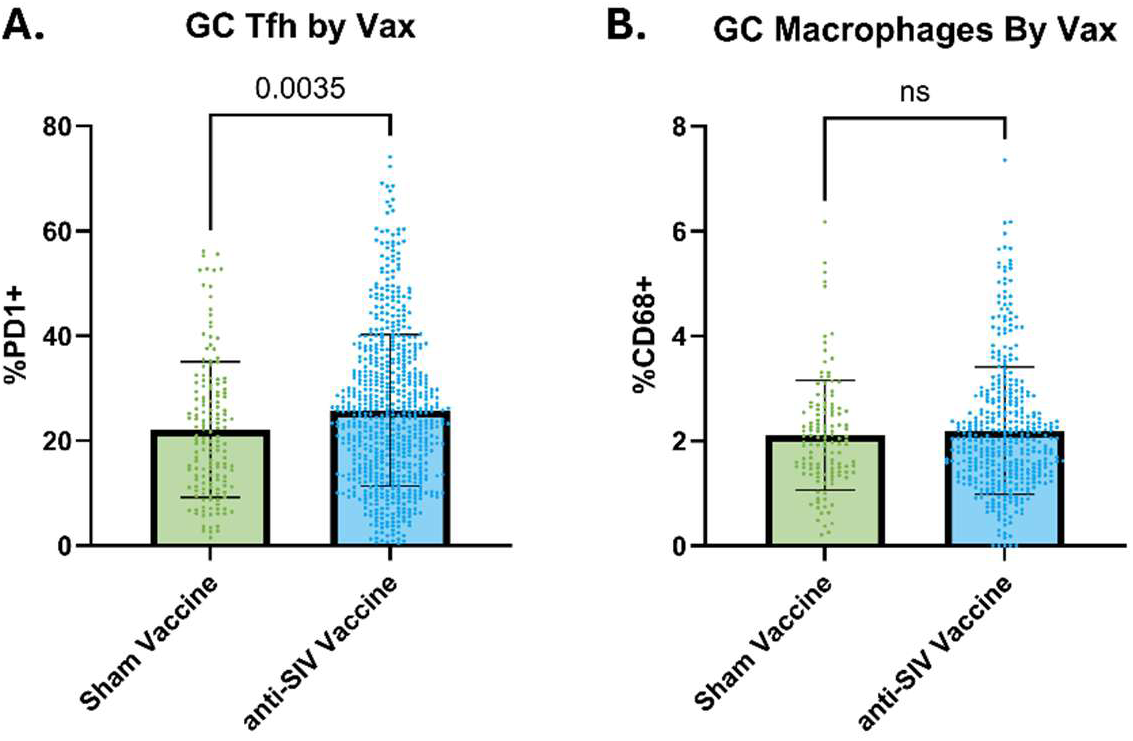
Anti-SIV T cell-directed vaccine increases GC Tfh cells in a macaque model of SIV infection. **A.** Analysis of NHP GC Tfh cells between the sham vaccine group (green, n=2 NHP) and the group vaccinated with a heterologous adenovirus vaccine encoding Tat, Vif, Rev, and Vpr (blue n=6 NHP). Each dot represents a single GC, data represents the compilation of multiple tissues (mediastinal lymph node, axillary lymph node, mesenteric lymph node, and spleen). **B.** Analysis of GC Macrophages between the sham vaccine group (green, n=2 NHP) and the anti-SIV vaccinated group (blue n=6 NHP). Each dot represents a single GC, data represents the compilation of multiple tissues (mediastinal lymph node, axillary lymph node, mesenteric lymph node, and spleen). Note: ns = p>0.05.

## Discussion

The germinal center is not only an invaluable component of the adaptive immune system but also a main viral tissue reservoir in PLWH. Tfh cells are known to be highly HIV-permissive and key sites of viral replication, yet the impact of HIV on GC Tfh cells and GC composition and morphology have been poorly characterized (4, 9–12). The data presented here were possible due to access to rare clinical samples from PLWH. We leveraged the insights provided by the clinical samples to ask specific questions using complementary models as part of a new approach methodology strategy combining clinical samples, non-human primate samples, and organoid and explant models. Using ex vivo HIV-exposed tonsil explants to explore Tfh transcriptomic changes, we demonstrated decreased Tfh support for B cells and changes in Tfh adhesion and trafficking receptors consistent with the findings we obtained from samples from PLWH. In the in vitro HIV-infected lymphoid organoids, we observed that some strains of HIV rapidly cause Tfh depletion and may disrupt GC-like structure formation. Finally, in an NHP model, we determined that GC Tfh cells can be protected by vaccination prior to SIV exposure, offering a potential proof of concept that adaptive immune changes in PLWH with suppressed virus could be improved.

The changes seen in GC Tfh cells in response to HIV noted in this study build substantially on a small body of literature that has been limited by access to incredibly rare tissue samples and has a broad range of implications in PLWH and potentially in PLWOH as well. The 33% decrease in GC Tfh cell proportion and reduced density in PLWH would be expected to disrupt the Tfh-B cell signaling that is essential for B cell survival and maturation (14, 27). Additionally, Tfh cells are one of the major sources of CXCL13, which interacts with the CXCR5 receptor to stimulate germinal center formation and maintenance (14). Tfh cells within the GC are known to polarize to the light zone of the germinal center and thus should be relatively dense and closely positioned. Our data showing decreased density and increased distance between Tfh cells are consistent with light zone dysfunction. Losing Tfh support in the light zone may help explain why PLWH exhibit increased morbidity and mortality from vaccine-preventable diseases and why vaccine responses are less durable in PLWH than PLWOH (28). Notably, our data demonstrate that GC Tfh cells also seem to decrease with increased age in PLWH, although these findings were not statistically significant in these data. This GC Tfh loss driven by HIV might combine with aging to provide two hits to the immune system that further explain the demonstrated immunological challenges in aging PLWH with suppressed viral load and robust CD4 counts (29). Decreased GC size in PLWH, as observed here, might also contribute to decreased B cell responses and antibody functionality. GC hyperplasia is a typical response to pathogens and facilitates high-affinity antibody production, which implies that smaller GCs may have reduced functionality and responses to pathogens and/or vaccinations (8). Previous studies of GCs in PLWH, in the early days of antiretroviral therapy, have demonstrated GC hypertrophy during early disease, with shape eccentricity and GC atrophy as acquired immunodeficiency syndrome (AIDS) began (30–32). Our data indicate a decrease in GC size in a predominantly ART-treated cohort, consistent with immune hyporesponsiveness. GC size also decreases with age in both PLWOH and PLWH, indicating that part of immunosenescence may be derived from changing GC dynamics.

Given the inherent limitations of human clinical trials to experimental modification and of ex vivo models to recapitulate full in vivo physiology, NHP models can provide a valuable mechanism to study the impacts of potential therapies and prevention strategies on Tfh cells. In an NHP model of SIV infection, untreated SIVmac251 causes a decrease in Tfh cells in the spleen (21). Here, we demonstrate that a T cell-directed SIV vaccine is associated with a small but significant increase in GC Tfh cells in GCs compiled from multiple immune tissues including the spleen, preventing the decrease observed in prior studies. Further work is necessary to determine the immunological relevance of this change and its magnitude over longer time courses; however, this suggests that vaccination and potentially other immune-mediated therapies could provide a protective benefit to GC Tfh cells.

Our data also demonstrate that other GC cell types are changing in PLWH, namely CD8+ T cells and macrophages. Classically, CD8 T cells, the main cytotoxic effectors, are largely excluded from or dysfunctional within GCs, which is thought to underlie HIV reservoir persistence (33–35). It is well documented that at the blood level, CD8 T cells are dysfunctional in PLWH early on in infection (36). Although relatively rare within GCs, CD8 T cell recruitment and function within GCs are thought to play a key role in the elite controller phenotype, in which PLWH control viremia without ART and display significantly delayed or nonexistent progression to immunodeficiency (36–39). Our study reinforces that these cytotoxic effectors, in line with previous reports, overcome their relative exclusion from GCs in PLWH (34, 37, 40). While other groups have also demonstrated increased recruitment of CD8 T cells to GCs in PLWH, the relationship between follicular CD8 T cell concentrations and disease severity remains unresolved (41–44). The strong correlation between GC CD8 T cell count and patient viral load supports the hypothesis that HIV drives GC CD8 T cell recruitment. Our quantification demonstrates a ∼6-fold increase in CD8 T cell proportion in GCs and a subsequent decrease in the distance between CD8 T cells and potential reservoir cells such as Tfh and macrophages that is greater than would be expected based on changes in cell number and GC size. This increased proximity could provide CD8 T cells with greater opportunity to mediate immune control of viral replication; however, this response is clearly insufficient or inhibited in some manner.

The decreased percentage of GC macrophages in PLWH that we observed was unexpected. A priori, we anticipated that increased B cell turnover due to chronic immune stimulation and reservoir cell death in HIV would increase the number of macrophages to clear cellular debris (45). The decreased GC size and thus decreased number of apoptotic B cells may account for the slight decrease in macrophages, or they may be targets of cytotoxicity mediated by the increased CD8 T cells.

The impact of HIV on the formation and function of GCs has been limited by the rarity of lymphoid-derived mononuclear cells from PLWH. We used an in vitro model of HIV infection in tonsil-derived organoids as a way to interrogate this phenotype. The organoid model is well described as a functioning adaptive immune system in vitro that forms GC-like structures and performs GC-like functions(46). Based on our organoid data, acute HIV infection does have an impact on GC-like structures in the organoid model. The trend toward decreased GC-like surface area in the HIV-infected wells, while not statistically significant, does follow what is observed in human tonsil tissue from PLWH (Figure 2). It may be that the 7-day organoid experimental timeline was insufficient to allow full GC-like structure remodeling driven by HIV. It is interesting to note that the trend toward decreased GC-like surface area and size diverges from previously published data demonstrating hypertrophy of GCs in acute HIV infection(31, 47).

This is likely explained by the in vitro nature of these organoids, which precludes the recruitment of external cells that could drive the hypertrophy observed in whole organisms (30). The depletion of Tfh cells from the X4-tropic virus-infected organoids is dramatic, poignantly highlighting HIV’s capacity to cause Tfh cell depletion. The R5-tropic ADA virus, meanwhile, does not seem to impact Tfh numbers and yet has a similar trend toward decreased GC-like surface area, raising the possibility that it may still be negatively impacting Tfh function. While this is far from a complete mechanism, it is demonstrable that HIV alters GC dynamics in both acute and chronic settings in a manner that implies Tfh dysfunction or absence.

Imaging-based technologies provide incredibly powerful approaches for generating data on localization and spatial relationships between cell types, but unfortunately provide little in the way of mechanisms. To extend our analysis of this phenotype beyond evaluations of germinal center size, shape, and cellular composition, scRNA sequencing was used to interrogate the potential mechanisms underlying decreases in GC Tfh cells and GC alterations more broadly. The analysis results identified decreased enrichment of pathways supporting B cells in HIV-exposed Tfh cells. This represents a decrease in Tfh support for B cells very early following HIV exposure, which supports a previous finding describing early changes in blood-derived immune cells from PLWH (36). Decreased Tfh support for B cells decreases B cell survival and could potentially explain the decreased GC size in PLWH (27). The GO analysis also provides insights into potential mechanisms behind the decrease in GC Tfh cells in PLWH. Numerous genes and pathways associated with immune cell trafficking and adhesion are less enriched in the HIV-exposed tonsil explants relative to unexposed controls (23, 48–50). It is also noteworthy to compare previous flow cytometry-based findings of an increase in Tfh cells in PLWH lymph nodes with our findings of a decrease in GC Tfh cells (9, 19). Based on the RNA sequencing data, it is plausible that HIV may inhibit the ability of Tfh cells to migrate or be retained at specific sites such as GCs, causing the localized, GC-specific decrease in Tfh cells observed here. We intend to explore this theory further in future studies through techniques such as spatial transcriptomics.

We acknowledge that there are limitations to the work reported here. Chiefly, although this sample set is relatively robust given the rarity of these samples, it remains a small sample size that limits our ability to extrapolate from the data presented. Due to the scarcity of tissue and the limitations of working with FFPE tissue, many of the desired experiments on PLWH-derived tissue are unable to be performed. Additionally, we acknowledge that the ex vivo tonsil explant acute HIV infection model used in the scRNA sequencing experiments, the in vitro lymphoid organoids, and the chronic natural infections represented in human tonsil samples are not directly comparable. However, this combination of approaches was necessary to gain insight into the tissue-based HIV reservoir and is consistent with cross-validation approaches to interrogate complicated phenotypes. We believe each of these models can provide translatable insights and data that are relevant to natural infections and intend to explore them further in future studies.

Lastly, we recognize that there are differences between lymphoid organs and that the mucosal surfaces of tonsils and exposure to the oropharyngeal space may impact their function. However, tonsils remain an accessible lymphoid tissue that does not require medically unnecessary biopsies or surgeries to access, supporting the ability to obtain them from a larger subset of participants with more generalizable demographics. Moreover, mucosal sites are a frequent point of initial infection in HIV and are thus well suited to serve as a model in this context.

## Conclusion

Studies of the tissue microenvironment in HIV have historically been limited by access to the requisite tissue. Here, we leveraged our access to rare clinical specimens of lymphoid tissue from PLWH to survey the impact of HIV on germinal center form and function. To complement this analysis, we applied in vitro infection of both explant and organoid models with HIV, as well as analysis of data from NHP samples, to assess the impact of vaccine intervention on the phenotypes identified here. The present study demonstrates that the GC microenvironment is substantially altered in people living with HIV. The decrease in Tfh cells, decreased GC size, and increased presence of CD8 T cells all likely negatively impact GC function. GC size also decreases with age and may put PLWH at additional risk of immunodeficiency as they age despite antiretroviral therapy. HIV’s negative impacts on Tfh ability to support B cells and decreased expression of genes associated with cell trafficking and adhesion may be driving the changes observed in GCs. While the decrease in Tfh cells in PLWH is likely to have a negative impact on immune function, our data demonstrate that anti-SIV vaccination increases GC Tfh cells in an NHP model of infection, suggesting that these changes may be reversible or treatable. The root causes underlying immune dysfunction in PLWH on ART remain a complex issue, but this study provides new evidence that HIV specifically alters germinal centers, which may contribute to weakened immunity in PLWH.

## Materials and Methods

Sample Collection: All TGH samples were collected in accordance with IRB-approved protocols. Patients recommended for tonsillectomy by their attending ENT physician were consented by a study team member prior to surgery (Table 1). Each tonsil was divided in half, with one half placed in 10% neutral buffered formalin for approximately 48 hours for fixation and the other processed fresh for mononuclear cell isolation as previously described(46). Following fixation, tissue was dehydrated and paraffin-embedded for downstream analysis. PLWH tonsil samples were collected similarly and provided through collaborations with investigators at the Africa Health Research Institute (AHRI) and University of KwaZulu-Natal (UKZN).

**Table 1.** Baseline demographic and HIV-related clinical characteristics of study participants.

| PLWH |  | N =21* |
| --- | --- | --- |
| *South African Cohort, n=21* |  |  |
| Age, years |  | 28.7 (12.0) |
| Age range, years |  | 2 – 48 |
| Sex | Male | 2 (9.5) |
|  | Female | 19 (90.5) |
| CD4 count, N=19<sup>†</sup> | cells/μL | 490.9 (51.0-876.0) |
| Viral load, N=15 | <20 copies/mL | 9 (60.0) |
|  | ≥20 copies/mL | 6 (40.0) |
| Viral load among quantifiable results, N=6<sup>‡</sup> | copies/mL | 42,068 (120-110,000) |
| PLWOH |  | N=40* |
| *South African Cohort, n=6* |  |  |
| Age, years |  | 31.8 (9.9) |
| Age range, years |  | 19 – 44 |
| Sex | Male | 5 (83.3) |
|  | Female | 1 (16.7) |
| *Tampa Cohort, n=34* |  |  |
| Age, years |  | 14.7 (11.5) |
| Age range, years |  | 3 – 39 |
| Sex | Male | 15 (44.1) |
|  | Female | 19 (55.9) |

\*Values are presented as n (%), mean (SD), or median (IQR), as appropriate.
†Median (IQR) for CD4 count was calculated among participants with an available CD4 count.
‡ Median (IQR) for viral load was calculated among patients with a quantifiable viral load ( $\geq 20$ copies/mL).

Immunohistochemistry (IHC): Formalin-fixed, paraffin-embedded tonsils were sectioned at 4–5 µm using a microtome. Slides were deparaffinized and rehydrated through xylene and graded ethanol solutions, followed by heat-induced epitope retrieval in a citric acid-based solution using a decloaking chamber. Slides were blocked with 2% milk for a minimum of 1 hour, followed by peroxide and protein blocking according to the manufacturer’s protocol (Epredia DAB Quanto Detection System). Primary antibodies were diluted as indicated in Table 2 and incubated on tissue sections for a minimum of 1 hour at room temperature. Detection was performed according to the manufacturer’s protocol. Slides were hematoxylin counterstained, mounted, and coverslipped. Representative images for each stain are shown in Supplemental Fig 2.

**Table 2:**
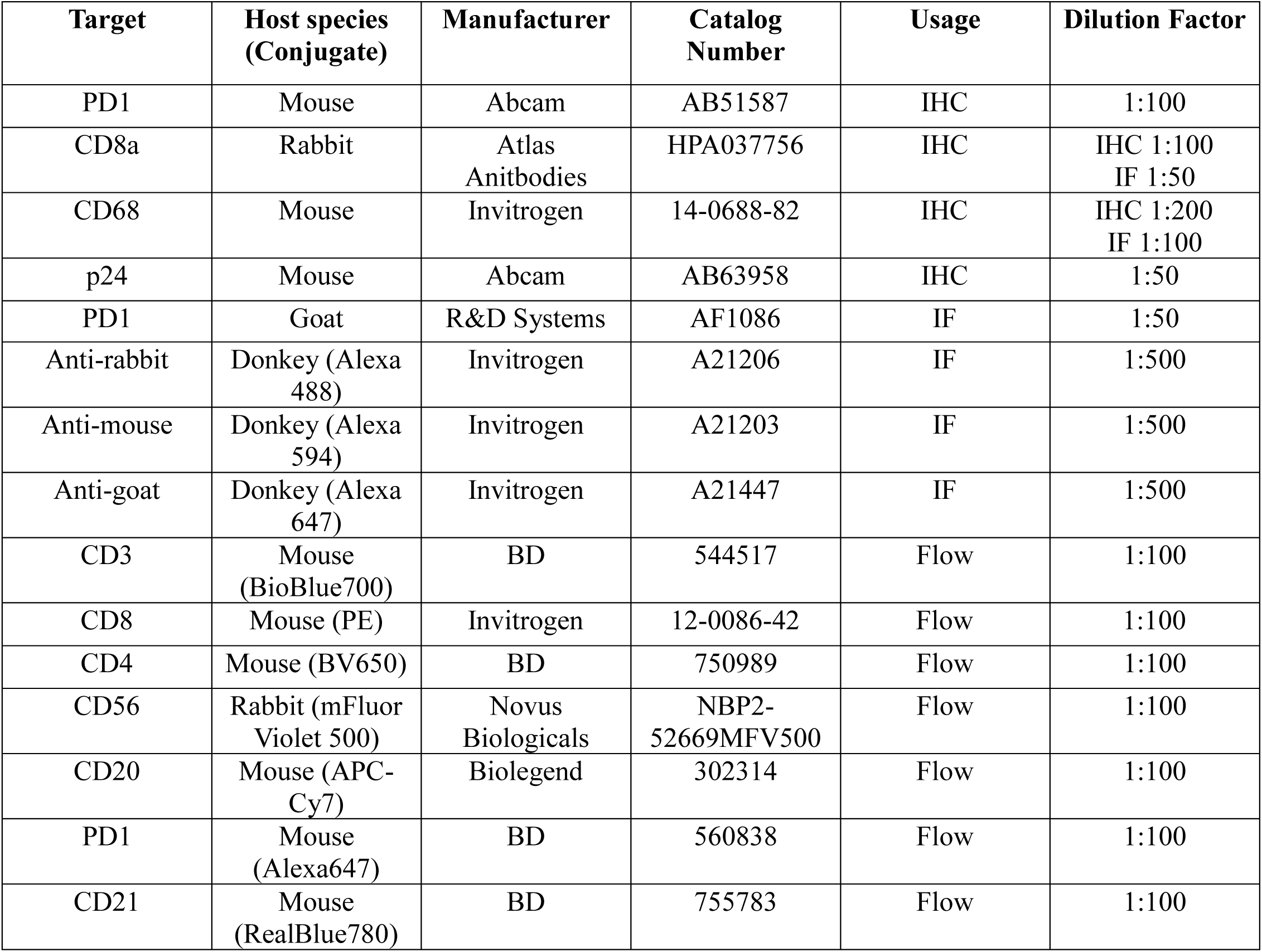
Details of antibodies used in these experiments. All antibodies were diluted in blocking solution.

| Target | Host species (Conjugate) | Manufacturer | Catalog Number | Usage | Dilution Factor |
| --- | --- | --- | --- | --- | --- |
| PD1 | Mouse | Abcam | AB51587 | IHC | 1:100 |
| CD8a | Rabbit | Atlas Anitbodies | HPA037756 | IHC | IHC 1:100<br>IF 1:50 |
| CD68 | Mouse | Invitrogen | 14-0688-82 | IHC | IHC 1:200<br>IF 1:100 |
| p24 | Mouse | Abcam | AB63958 | IHC | 1:50 |
| PD1 | Goat | R&D Systems | AF1086 | IF | 1:50 |
| Anti-rabbit | Donkey (Alexa 488) | Invitrogen | A21206 | IF | 1:500 |
| Anti-mouse | Donkey (Alexa 594) | Invitrogen | A21203 | IF | 1:500 |
| Anti-goat | Donkey (Alexa 647) | Invitrogen | A21447 | IF | 1:500 |
| CD3 | Mouse (BioBlue700) | BD | 544517 | Flow | 1:100 |
| CD8 | Mouse (PE) | Invitrogen | 12-0086-42 | Flow | 1:100 |
| CD4 | Mouse (BV650) | BD | 750989 | Flow | 1:100 |
| CD56 | Rabbit (mFluor Violet 500) | Novus Biologicals | NBP2-52669MFV500 | Flow | 1:100 |
| CD20 | Mouse (APC-Cy7) | Biolegend | 302314 | Flow | 1:100 |
| PD1 | Mouse (Alexa647) | BD | 560838 | Flow | 1:100 |
| CD21 | Mouse (RealBlue780) | BD | 755783 | Flow | 1:100 |

Immunofluorescence (IF): The immunofluorescence protocol was identical to the IHC protocol through primary antibody incubation. IF primary antibodies were incubated overnight at 4°C. Secondary antibodies were prepared as indicated in Table 2 and incubated for 1 hour at room temperature in the dark. Slides were mounted with DAPI-containing Fluoromount-G medium and imaged.

Imaging: All images were acquired using an Echo Revolution Microscope with a 40× objective. Exposure times were automatically determined by the instrument software. Focus mapping points were used to ensure images were captured in optimal focus across the tissue. IF images were acquired using DAPI, FITC, Texas Red, and Cy3 filter sets. All IHC and IF images were exported as TIFF files and uploaded to Box for downstream analysis.

Image Analysis: All images were analyzed using QuPath-0.5.1 software(51). For IHC images, germinal centers were manually identified and annotated. Only in-focus, non-torn or folded, well-stained, high-quality germinal centers were included in the analysis. A pixel classifier was trained for each image using manually identified DAB-stained and unstained cells to account for variability in staining intensity between images. The classifier was then used in combination with the cell detection feature (Supplemental Fig 3). For p24 staining, cells were quantified using the Positive Cell Detection feature, with a single threshold applied across all images. Tfh-Tfh distances were calculated using a modified version of the Detection Centroid Distances 2D function. A custom script was used to prevent cells from identifying themselves as the nearest Tfh cell. Annotation and detection measurements were exported to GraphPad Prism for statistical analysis.

IF images were analyzed similarly using QuPath. GCs were manually identified and annotated, and cells were detected using the Cell Detection function on the DAPI channel. Fluorescently labeled cells were identified using a measurement classifier, with the threshold manually adjusted for each fluorophore to maximize detection of positive cells while minimizing classification of cells labeled in other channels. Cell-cell distances were calculated using the Detection Centroid Distances 2D feature. Cells/detections labeled in multiple channels (e.g., CD8:Tfh:Mac) were excluded from analysis to minimize the effects of autofluorescence and nonspecific secondary antibody binding. Representative images of each channel and analysis step are shown in Supplemental Fig 3.

Single-cell RNA (scRNA) sequencing Analysis: This analysis was conducted using the publicly available, previously published dataset GSE232477 (26). Briefly, these data were collected from an experiment using HIV NL-CI- exposed tonsil explants and naïve controls. All analyses were performed using RStudio 2026.01.2. Cells from Cell Ranger outputs were filtered based on the number of genes detected and the percentage of mitochondrial genes, and doublets were removed using scDblFinder (v1.24.10) (52). Due to large interpatient variation across samples, samples were integrated using the Harmony integration method in Seurat (v5.4.0) (53). Cell types were annotated based on canonical cell-type markers. T cells were subsetted from the whole tonsil dataset and reclustered to annotate T-cell subpopulations. Differential gene expression analysis was conducted between HIV-exposed and HIV-naïve Tfh cells. Gene ontology (GO) term analysis was performed on genes differentially expressed between exposed and unexposed Tfh cells using ClusterProfiler (v4.18.4).

Lymphoid Organoid Model: The tonsil mononuclear cell (TMC)-derived organoid model was adapted from Wagar et al. 2021 (46). TMCs were isolated from freshly harvested tonsils using a Ficoll gradient, frozen, and stored in liquid nitrogen until organoid formation. TMCs were thawed and plated in 48-well plates, followed immediately by HIV exposure (Fig 4A). Plasmids encoding for HIV stocks were obtained from the AIDS Reagent Program and expanded in-house. Experiments used HIV NL4-3 and HIV NL4-3 ΔADA-GFP to model CXCR4- and CCR5-tropic infection, respectively. Organoids were infected at the EC50 concentration determined using a TZM-bl luciferase assay. GC-like cell aggregates began forming spontaneously within 24 hours of plating (Fig 4B). Organoids were imaged on days 1, 4, and 7, and media were exchanged on day 4 following imaging. Organoid images were analyzed using QuPath-0.5.1. An identical circular region in the center of each well was analyzed for each image using a pixel classifier trained separately for each image.

Flow Cytometry: All wells were harvested on day 7 following imaging and stained for flow cytometry. Following staining, cells were fixed using BD Fix/Perm and stored overnight before acquisition on a Miltenyi MaxQuant16 instrument. Gates were established using fluorescence-minus-one (FMO) controls. Tfh cells were defined as CD4+PD1+ cells. All flow cytometry analyses were performed using FlowJo version 10.10.1.

Total IgG ELISA: Organoid media were collected on days 7 and 10 for measurement of total IgG by ELISA. The ELISA was performed according to the manufacturer’s protocol (Thermo Fisher Catalog number BMS2091). Absorbance was measured at 450 nm using a Varioskan ALF instrument.

Primate Study: The non-human primate (NHP) experiment was conducted by collaborators at the Tulane National Biomedical Research Center as part of a study of T cell-directed HIV vaccination strategies in rhesus macaques. Eight NHPs were included: six received an adenoviral vaccine containing HIV *tat, vif, rev,* and *vpr* (hAd19a/64/CMV/Ii/tat-vif-rev-vpr), while two received a sham vaccine containing an empty viral vector (hAd19a/64/CMV). Twenty-five weeks later, all animals received rectal SIVmac251 exposure once weekly for 10 weeks. Viral loads were monitored throughout the study. Following detection of viremia in all animals, ART was initiated, and animals were sacrificed at week 63. IHC analysis was performed on mediastinal, axillary, and mesenteric lymph nodes and spleens obtained at necropsy using the same methodology as for human tissues.

Statistical Analysis: All statistical analyses were performed using GraphPad Prism 10.6.0. Pairwise comparisons were performed using two-tailed Welch’s *t*-test or the Mann-Whitney U test, as appropriate. Multiple comparisons were performed using one-way ANOVA with Tukey’s multiple-comparisons test. Associations between continuous variables were evaluated using linear regression. An α threshold of 0.05 was used for all analyses.

## Acknowledgements

We would like to acknowledge our international collaborators at AHRI, UKZN, and Tampa General Hospital without whom this study would not have been possible. We would also like to acknowledge our patients who generously donated their tissues for this study. Funding for this study was provided by the Pediatric Adolescent Virus Elimination (PAVE) Martin Delaney Collaboratory UM1AI164566 and by the Tulane National Biomedical Research Center grant P51OD011104. We thank BEI Resources, NIAID, NIH for providing the reagents used in this study. Lastly, our thanks go to Pyone Pyone, Aye, Faith Schiro, and the Unit of Collaborative Research (RRID SCR_024610) for providing the NHP tissues, and to the AIDS reagent program for their contributions to key cell lines and reagents.

---

**S1 Fig. HIV-mediated changes on germinal center Tfh cell numbers do not correlate with patient viral load, and macrophage counts do not vary with age, CD4 count, or viral load. A.** PLWH correlation of GC Tfh cells with patient viral load, n = 14. **B.** GC macrophage correlation with age in PLWOH, n = 12. **C.** GC macrophage correlation with PLWH CD4 count, n = 11. **D.** GC macrophage correlation with PLWH viral load, n = 10.

**S2 Fig. Representative images for IHC stains and example of QuPath IHC cell counting analysis. A.** Representative IHC stains for each analysis used in this study, each image contains an entire germinal center. Stains are from left to right: PD1, CD8a, CD68, and p24. **B.** A breakdown of the individual steps used in the IHC image analysis. From left to right: manual identification of GC, green transformation of all pixels identified as DAB stained following pixel classifier training, results of hematoxylin based cell detection, and lastly the combined pixel classification and cell counter which identifies Tfh cells and negative cells.

**S3 Fig. Representative images of IF stains and example of QuPath spatial analysis. A.** Representative images of each stain alone or an individual stain + DAPI, and a representative image of a GC with all stains. **B.** A breakdown of the individual steps used in the spatial analysis. From top left to bottom right: Manual identification of the GC, results of cell counting using the DAPI channel, results of identified cells after classification using the object classifier, and a higher magnification image of the cell-cell distance lines used in the IF spatial analysis.

**S4 Fig. Additional figures generated during scRNAseq analysis. A.** UMAP of all explant cells from all donors post-integration. **B.** UMAP of all explant cells with cell annotations assigned by canonical markers. **C.** UMAP of all T cells split by HIV exposure.

**S5 Fig. HIV exposure does not cause changes in GC-like surface area in tonsil organoids at early time points. A.** Total GC-like surface area in tonsil organoids on day 1 of culture, n = 4 donors, ns = p>0.05. **B.** Total GC-like surface area in tonsil organoids on day 4 of culture, n = 4 donors, ns = p>0.05. **C.** ELISA results of organoid culture media assessing total IgG production of day 7 organoids. n = 8 donors. ns = p>0.05.

